# Learning Discriminative Representations of Superimposed P waves With Weakly-Supervised Temporal Contrastive Learning

**DOI:** 10.1101/2024.04.28.591427

**Authors:** Jakub Hejc, Richard Redina, David Pospisil, Ivana Rakova, Jana Kolarova, Zdenek Starek

## Abstract

Electrocardiography (ECG) wave morphology and timing provide critical information for diagnosing arrhythmias and conduction abnormalities, allowing risk stratification for various cardiac diseases. However, extraction of these features becomes challenging in the presence of superimposed waves from distinct cardiac chambers, a common occurrence during pathological rhythms. This work proposes a novel Surrogate-boosted Temporal Contrastive Representation Learning (S-TCRL) framework to address this challenge. S-TCRL leverages weak labels, readily obtainable from invasive catheter examinations, to extract latent representations of superimposed P waves.

We reformulate the problem from object-wise to sample-wise incomplete information by employing surrogate labels. A 1D fully-convolutional feature pyramid network (FPN) extracts multi-scale features from ECG signals. These features are segmented into equal-sized temporal regions, whose labels are inferred from individual samples using a multiple-instance learning (MIL) paradigm. Non-sequential embeddings are generated to facilitate alignment-free cosine similarity estimation. A temperature-scaled cross-entropy loss function minimizes the distance between embeddings of similar regions (likely containing P waves) while maximizing the distance between dissimilar ones.

The framework’s efficacy is evaluated on a custom ECG dataset comprising 3265 short-term recordings from 708 individuals undergoing catheter ablation. S-TCRL achieves significant improvement in the downstream P wave segmentation task compared to two baseline MIL methods. The average recall and precision for both P wave boundaries reach 70.0% and 80.0%, respectively, exceeding the baselines’ 63.5% and 67.5%. The results demonstrate the potential of S-TCRL for embedding representation of superimposed P waves and its generalizability to tasks such as arrhythmia classification.

## Introduction

Accurate segmentation and characterization of P waves in electrocardiograms (ECGs) are crucial for diagnosing cardiac arrhythmias and studying atrial electrophysiology. P wave morphology and timing provide valuable clinical markers for various conditions (Kimura-Medorima et al., 2018), including atrioventricular (AV) blocks, supraventricular tachycardias (SVTs), and atrial fibrillation (AF) (Brady et al., 2017; Stafford et al., 1991). These distinctions are essential for guiding therapy and patient prognosis.

Differentiating arrhythmias based on P wave characteristics can be challenging. For instance, complete atrio-ventricular dissociation, characterized by independently triggered activity within atria and ventricles, is a hallmark of ventricular tachycardia (Brugada et al., 1991; Brady et al., 2017). In contrast, SVTs with wide QRS complexes often exhibit varying degrees of temporal coupling between the P wave and the QRS complex. P wave frequency variability can aid in differentiating focal atrial tachycardia from isthmus-dependent atrial flutter (Roberts-Thomson et al., 2006). Conversely, the absence of P waves in an ECG recording is a characteristic finding in atrial fibrillation. Accurate distinction between paroxysmal atrial fibrillation (AF) and premature atrial contractions (PACs) or other rhythm disorders is critical for quantitatively assessing the temporal burden of atrial fibrillation, a risk factor for stroke, heart failure, and mortality (Chen et al., 2018).

Extracting P wave features becomes particularly difficult in the presence of abnormal rhythms due to several factors (Hernández et al., 2000; Portet, 2008): (a) low atrial signal energy and poor signal-to-noise ratio (SNR), (b) dominant frequency components (5-30 Hz) overlapping with other waves (Tereshchenko et al., 2015), (c) wave superimposition during abnormal P wave timing, (d) high inter-individual variability in morphology, especially in complex arrhythmias and the presence of structural changes, and (e) the scarcity of data with reliably labeled P wave positions for training advanced machine learning models.

This work introduces a surrogate-boosted temporal contrastive representation learning (S-TCRL) framework to extract latent representations of superimposed P waves in a weakly-supervised manner. S-TCRL leverages the unique opportunities in cardiac electrophysiology, where auxiliary information is often obtainable from other clinical measurements. For instance, during invasive electrophysiology procedures (EPs) in arrhythmia patients, a surrogate dataset combining surface ECGs and intracardiac signals can be acquired to simplify labeling.

Our previous work suggests that directly modeling the probability distribution of P waves might not be optimal for representation learning. S-TCRL addresses this by encoding the P wave into a compact embedded space via non-linear mappings that preserve the similarity between specific temporal regions of the signal. This approach facilitates the extraction of microfeatures by deep neural networks, potentially leading to improved arrhythmia classification and a better understanding of model decision-making processes.

Automated extraction of individual atrial features from ECG recordings with superimposed waves has the potential to: (a) enhance the diagnostic yield of ECGs, leading to more accurate arrhythmia identification, (b) reduce the economic burden associated with clinical ECG interpretation, particularly in long-term monitoring scenarios; (c) enable the development of inherently interpretable deep learning models for downstream tasks.

## Related works

### P wave segmentation algorithms

P-wave detection algorithms can be broadly categorized into three main approaches. Local search methods leverage the correlation between the P-wave and chosen base functions to enhance atrial activity. These functions are used to extract features like extrema and inflection points within the TQ segment of the ECG signal for P-wave detection. Common choices for feature extraction include wavelet transform (Martinez et al., 2004; Li et al., 1995; Sahambi et al., 1997), quadratic splines Martinez et al. (2004); Li et al. (1995), and derivatives of Gaussians (Martinez et al., 2004). The simplicity of these methods has driven research efforts towards optimizing parameters affecting base function shape (Spicher and Kukuk, 2020; Lenis et al., 2016), thresholding rules (Lenis et al., 2016), and computational efficiency (Di Marco and Chiari, 2011).

McSharry et al. (2003) proposed a nonlinear parametric model based on differential equations to generate synthetic ECG data. This model was subsequently expanded by Akhbari et al. (2016) and Hesar and Mohebbi (2019) using additional parameters and a Kalman filter-based optimization to facilitate P-wave position, width, and amplitude regression.

Other approaches explore nonparametric and nonlinear ECG transformations into latent spaces potentially correlated with the P-wave shape. These transformations include empirical mode decomposition (Hossain et al., 2019), phase transformation (Martínez et al., 2010; Saclova et al., 2022), and optimization using evolutionary algorithms (Panigrahy and Sahu, 2018).

Ventricular activity suppression methods aim to isolate the P-wave by removing ventricular activity from the ECG signal. A straightforward approach involves directly subtracting a template QRS-T complex from the original signal (Stridh and Sommo, 2001; Sippensgroenewegen et al., 2001; Marenco et al., 2003). This method requires R-wave positions for accurate template localization and often utilizes dynamic time warping to adapt the template shape to account for respiration and heart rate variations. Rieta et al. (2004) and Vaya et al. (2007) proposed blind source separation using linear independent component analysis to extract atrial activity from 12-lead ECG recordings, assuming statistical independence with non-Gaussian distribution between atrial and ventricular activity.

Deep learning methods have shown potential in ECG analysis tasks, but existing P-wave segmentation algorithms often adapt successful architectures from computer vision without addressing limitations like poor generalization on ECGs with pathological rhythms (Saclova et al., 2022; Portet, 2008). Examples include works by Costandy et al. (2020) and Duraj et al. (2022) who utilized the U-Net architecture with 2D and 1D convolutional kernels, respectively. Alternatively, recurrent neural networks such as long short-term memory (LSTM) networks, have been proposed to capture signal evolution over time (Qi et al., 2019; Peimankar and Puthusserypady, 2019; Malali et al., 2020).

### Contrastive representation learning on time series

Contrastive representation learning (CRL) offers a powerful paradigm for learning data embeddings where similar instances are brought closer and dissimilar ones are pushed apart. Initially established as a self-supervised technique (van den Oord et al., 2019; Hadsell et al., 2006; Gutmann and Hyvärinen, 2010; Chen et al., 2020), CRL leverages data augmentation to construct positive and negative instance pairs. Subsequently, various contrastive losses such as Triplet Loss (Schroff et al., 2015), Noise Contrastive Loss (van den Oord et al., 2019), and Normalized Scaled Cross Entropy Loss (Chen et al., 2020) are employed to maximize similarity between positive pairs while minimizing those of the negatives.

Recent advancements have witnessed the successful application of self-supervised temporal CRL (TCRL) in capturing evolving contextual representations within video sequences (Dave et al., 2022; Sermanet et al., 2017), EEG (Yuan et al., 2019; Hyvarinen and Morioka, 2016; Banville et al., 2021), and ECG signals (Kiyasseh et al., 2021; Mehari and Strodthoff, 2022; Sarkar and Etemad, 2022). These methods typically involve segmenting the sequence into fixed-length segments, from which positive and negative instances are sampled.

A pioneering work for a time series by Hyvarinen and Morioka (2016) proposed an unsupervised TCRL approach for feature extraction, assuming non-stationarity between temporal segments. The framework combined a deep neural network with logistic regression to predict a label for each segment based on its sequence order, enabling feature embedding extraction akin to nonlinear source separation.

In video sequence representation, Dave et al. (2022) introduced a temporal contrastive loss to learn temporally distinct features across video sequences. Here, self-supervised contrasting occurred across non-overlapping segments from a single video instance, as opposed to augmented variants of the same instance. Similarly, Sermanet et al. (2017) leveraged identical temporal segments from multi-view videos as positive pairs for a Triplet Loss, generating temporal embeddings for video frame classification and pose estimation.

A comprehensive review by Liu et al. (2023) delves deeper into self-supervised contrastive techniques for medical time series. In cardiac signal pretext modeling, Kiyasseh et al. (2021) employed a self-supervised contrastive approach for short-term ECG signals, maximizing similarity between segments from the same signal and across different ECG leads, resulting in multi-segment, multi-view coding. However, their work assumes that clinically relevant changes are unlikely to occur within short timeframes, which might not hold true for specific cardiac conditions. To address this, Mehari and Strodthoff (2022) investigated both conventional instance-based contrastive learning and LSTM-based predictive coding for self-supervised sequential feature extraction, achieving significant improvements in downstream arrhythmia classification. Sarkar and Etemad (2022) explored self-supervised pretraining with pseudo-labels for ECG-based emotion recognition. Their recognition network aimed to predict noise and morphology-aware augmentations applied to ECG signals, leading to subject-specific embeddings that enhanced recognition performance.

Banville et al. (2021) proposed a contrastive method for EEG signal feature embeddings based on relative positioning and segment shuffling. Their approach leveraged the assumption that appropriate data representations evolve slowly over time, implying that nearby time windows share a similar class, while distant windows likely stem from different EEG patterns. Additionally, temporal shuffling involved sampling two temporal segments as positive anchors, with a third segment drawn from nearby or faraway segments to create a triplet. A recognition network then predicted the temporal order of the triplet. In contrast, Cheng et al. (2020) focused on subject-aware representation learning by minimizing the difference between an original EEG sequence and its augmented duplicate, similar to van den Oord et al. (2019). More recently, Yuan et al. (2019) proposed an end-to-end model that jointly optimized a sparse autoencoder to capture temporal patterns on EEG and ECG sequences in the time-frequency domain. Contextual dependencies were modeled akin to word embedding techniques using a skip-gram model.

Current TCRL methods based on fixed-length segmentation suffer from limitations. Segmenting data can lead to information loss, particularly for long-range dependencies or variable-length patterns that span segments. This hinders comprehensive representation learning. Furthermore, choosing the optimal segmentation strategy can be challenging and computationally expensive. These limitations motivate the exploration of alternative TCRL techniques for more effective time series representation.

### Problem formulation

Extracting temporal features from ECGs to represent individual electrical sources, such as ECG waves, in a fully supervised manner assumes each sample has a complete reference annotation (label) in the training set. However, real-world signals rarely meet this assumption. Labels can be distorted by noise, incomplete, or entirely missing due to factors like the high cost of manual labeling, annotation errors, and interpretation ambiguity. Consequently, collected labels are often provided only for:

1. Subset of waves (object-wise incompleteness).
2. Some samples of each wave (sample-wise incompleteness).
3. A combination of both (A scenario frequently encountered in pathological ECGs).

To overcome object-wise label scarcity, we propose using surrogate data consisting of inexpensively acquired labels, such as those derived from intracardiac atrial signals from cardiac pacemakers or interventional electrophysiology procedures. This approach allows to construct a partially labeled set of waves *Y*, with their corresponding input data *X*.

We can formulate the problem as follows: Given an ECG time series represented by a set of instances *X* = {*x*_1_, …, *x*_*N*_} and a set of *M* distinct electrical sources *S* = {*s*_1_, …, *s*_*M*_}, where, in the simplest case, *M* = 1 for uniform atrial activation wave, we aim to determine whether source *m* was active at a specific time point *n*. The ground truth labels are unavailable for some instances in *X*. Therefore, the labeled data for *m*-th source is represented by *Y*_*m*_ = {*y*_*m*,1_, …, *y*_*m,K*_}, where *K ≪ N*. Here, *K* signifies the number of instances with known ground truth, and *N − K* epresents the number of unidentified instances of source *s*_*m*_.

The objective is to find a function composition *h ◦ z*: *X → Y*, where *h* learns temporal embeddings that differentiate between the original source waveforms based on relevant features, *Y* represents the incomplete ground truths, and *z* acts as an estimator that models the posterior probability *P* (*y*_*n*_ ∈ *s*_*m*_ | *X*_*τ*_) for both labeled and unlabeled instances. This estimation is based on the surrounding context captured within a local region of the time series *X*_*τ*_. The contextual relationship between samples and labels can be modeled using weakly supervised learning paradigms, such as Deep Multiple Instance Learning (Gonthier et al., 2022).

## Materials and Methods

### Data

3265 8-lead ECGs from 708 patients (41.7% female, median age 36.6 years, median duration 11.0 seconds) indicated for electrophysiology procedure were employed in this study. ECGs were recorded during electrophysiology examination using Abbott WorkMate 4.3 system at a sampling rate of 2000 Hz and a resolution of 78 nV/LSb. The acquisition unit included built-in hardware high-pass filter with a cutoff of 0.5 Hz and a notch filter at 50 Hz.

Individual signals were categorized into one of four groups based on visual evaluation of prevailing rhythm: a) sinus rhythms including premature beats, A-V blocks, and bundle branch blocks (SRs); b) supraventricular tachycardias (SVTs); c) atrial fibrillation and/or flutter (AF); and d) rhythm from artificial pacing (ST). Each category presents a varying level of segmentation difficulty. For instance, the SVT group predominantly includes P waves superimposed on ventricular activity, while the AF group mostly contains no localized atrial complexes. Overall, the datasets comprise of 1,771, 1,036, 458 and 770 annotated ECGs in SB, SVT, AF and ST group. Naive oversampling was applied to address the distribution imbalance between the groups.

### Surrogate labels

Electrophysiology study is crucial for investigating myocardial electrical properties and arrhythmia mechanisms (Katritsis et al., 2016; Issa et al., 2019). During the procedure, a multi-channel diagnostic catheter positioned within the coronary sinus (CS) is commonly used to aid in identifying the arrhythmia substrate. The recordings capture well-defined atrial activation patterns, representing a diagnostically important electrical component. While localizing atrial waveforms in pathological ECGs can be challenging or even impossible, intracardiac atrial complexes can be segmented relatively easily.

A surrogate set of inexpensive labels was obtained by segmenting multi-channel atrial electrograms using a custom-developed algorithm (Hejc et al., 2024). The algorithm identifies the earliest and latest atrial activation times across multiple electrode locations. These surrogate labels provide information about the propagation of the waveform across the left atrium, although details like position and duration remain incomplete. Figure 1 illustrates the steps involved in generating the surrogate label set *Y* ∈ {0, 1}.

**Figure 1.**
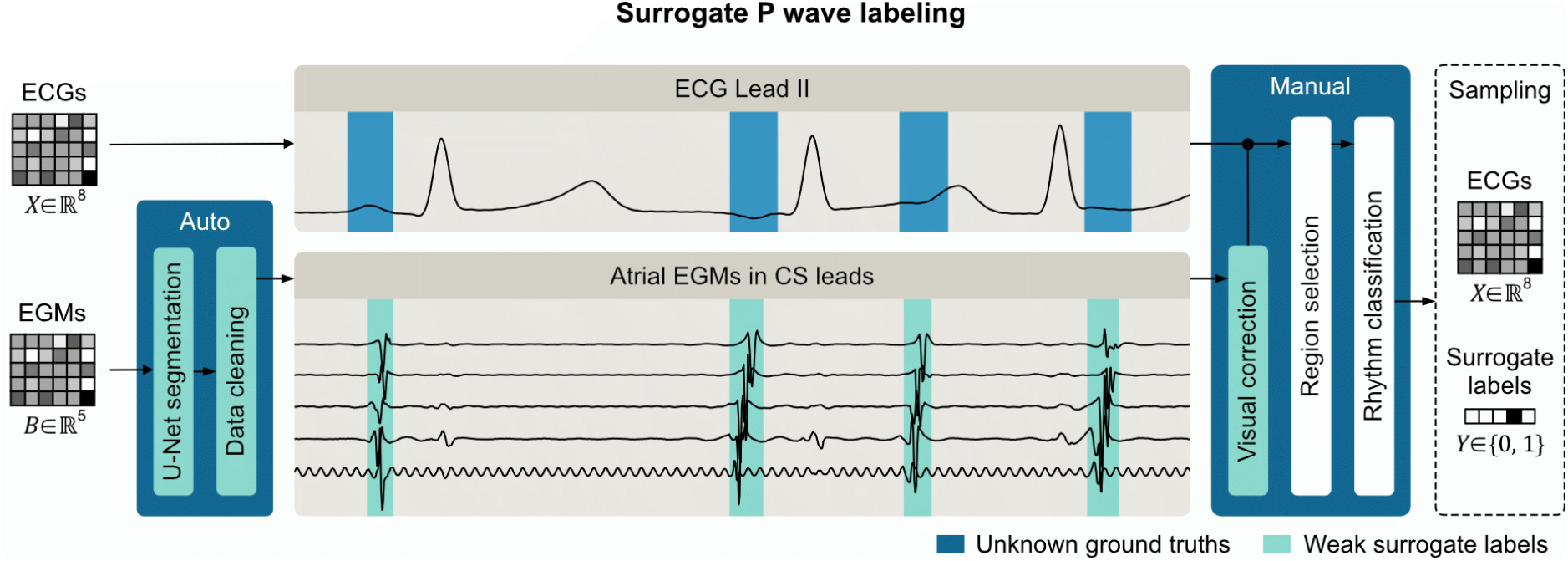
Schema representing the orchestration of 12-lead ECG signals and intracardiac EGMs during surrogate label preparation. Top panel depicts example of the ECG Lead II including expected ground truth segments (blue boxes) representing atrial activity occurrence and duration. Bottom panel depicts related EGM recorded at the same time with automatically segmented atrial beats (green boxes) forming a surrogate incomplete label of the atrial activity.

**Figure 2.**
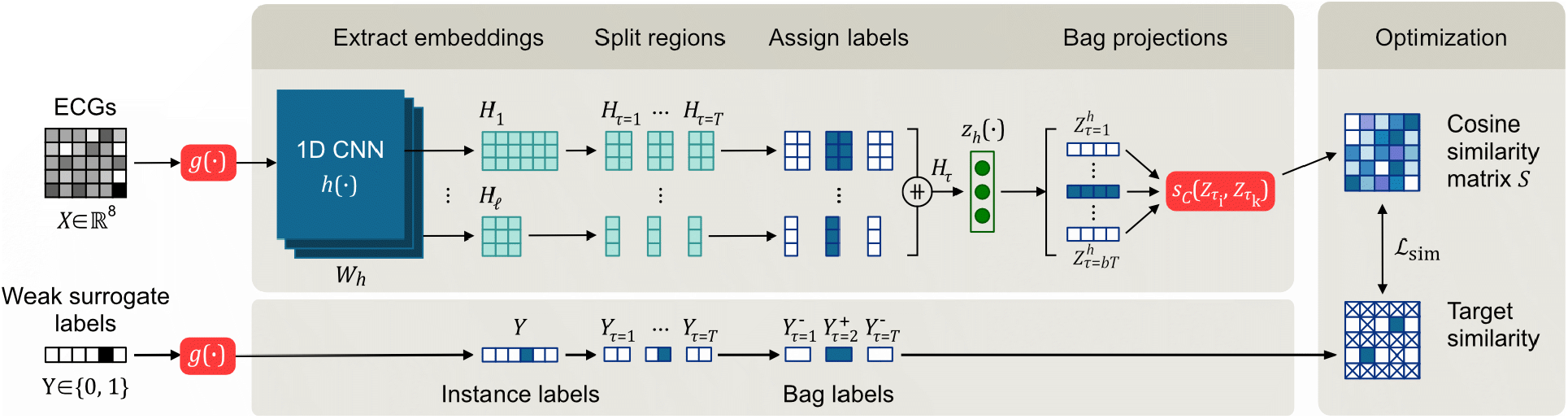
Surrogate-boosted temporal contrastive representation learning (S-TCRL) framework for atrial activity feature extraction based on intra-sequence and cross-sequence similarity. Functions *g, h, s*_*c*_, and *z*_*h*_ represent morphology and noise-aware augmentations, feature extractor based on deep convolutional neural network, cosine similarity function, and projecting network for modeling alignment-free similarity, respectively. Tensor *H* denotes deep temporal representation of the input multi-lead ECG signal *X*.

While irregular atrial fibrillatory waves were actively classified as a negative class due to their inherent temporal variability that spans the entire signal without a well-defined boundaries, they could potentially represent a distinct source within a multi-label framework for future investigations.

### Data partitioning

Stratified randomization (SR, SVT, AF and ST groups were used as strata) into 70:30 training (DS_T_) and validation sets (DS_V_) was employed to mitigate potential bias in quality metrics arising from the highly imbalanced representation of cardiac rhythms. The data were partitioned using patient-oriented approach ensuring non-overlapping patient selection in both DS_T_ and DS_V_. The optimal cross-set distribution was assessed using relative entropy.

### Preprocessing and standardization

ECGs were undersampled to 250 Hz using anti-aliased decimation. To remove potential baseline drift and extract the most relevant frequencies, ECGs were processed with a bandpass filter with a 0.5 and 40Hz cutoff frequency. ECGs exceeding ±12.436 mV were clipped to eliminate extreme voltage oscillations and then rescaled using z-score standardization.

### Morphology and noise aware augmentation

Simple affine transformations of ECGs are proposed to modulate underlying bipole directly representing waveform velocity increasing morphological variability. Morphological data augmentations (DAs) included sarcastically driven voltage scaling, polarity flipping, temporal translation, linear temporal scaling, and reversing the temporal axis. The latter method was employed to mitigate overfitting arising from the stochastic interrelationship within the P-QRS-T sequence.

A corrupted version of the original ECGs was generated by adding Gaussian white noise, simulated power line interference and by zeroing samples in one or more ECG channels. Noise power together with resulting signal-to-noise ratio (SNR) of corrupted ECG were stochastically modulated.

### Embeddings extraction

A temporal convolutional neural network (1D-CNN) handling ECG channels as a separate features (Vicar et al., 2020, 2021) was employed to learn a non-linear mapping *h*: *X → H*, where *h* is a feature extractor and *H* represents vector embeddings. The CNN incorporates a ResNet-50 backbone encompassing four residual stages with residual units combining preactivated 1 *×* 7 kernels and a 1 *×* 1 bottleneck with a 50% compression factor. The filter dimensions across encoder stages consist of 128, 256, 512, and 1024 filters. A feature pyramid network (FPN) (Lin et al., 2017) was integrated as a decoder allowing for the extraction of semantically rich multi-scale embeddings *H* = {*H*_1_, …, *H*_*ℓ*_}, where *ℓ* denotes number of decoding layers.

### Weakly supervised temporal contrastive representation learning

The objective was to find transformation *f*: *H → Ŷ* given the ground truths *Y*. Function *f* (*H*) is an estimator used to model the posterior probability *ŷ*_*i*_ ∈ *Ŷ* of being a part of the P wave for both labeled and unlabeled instances. The proposed method models *f* (*H*) using a combination of multiple instance learning (MIL) paradigm and augmentation-free time-contrastive learning representation (TCLR) to achieve temporaly informed deconvolution of atrial activity.

MIL frames the original problem to learning *f* (*H*) to predict the ground truth label *Y* ∈ {0, 1} of a bag given its instances. A combination of weakly supervised (Gonthier et al., 2022) and self-supervised strategies (Hénaff et al., 2020; Bachman et al., 2019) from image experiments was adapted to time-series embeddings by dividing *H* into equal-sized temporal regions, i.e. views or bag of local features, indexed by *τ* = {1, …, T}. A bag is defined as a subsequence *H*_*τ*_ *⊆ H*, and its label is inferred from individual instances using MIL paradigm. In other words, *H*_*τ*_ is considered positive if and only if at least one of its instances is labeled as positive.

Contrastive representation learning based on the method proposed by Chen et al. Chen et al. (2020) was employed as a pretraining technique to learn invariances in the problem space encompassing incomplete labels. Since true alignment between a pair 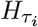 and 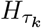 were not known, a projecting network *z*_*h*_(*H*_*τ*_), i. e. multilayer perceptron, was introduced in order to produce non-sequential embeddings 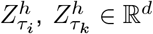. The alignment-free similarity can be then estimated using cosine similarity:

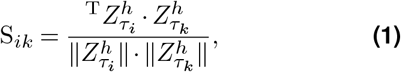

where 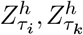represents the positive pair of metric embeddings derived from the two distinct regions of 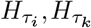 only if both contain at least a one instance from a positive class; all other pairings are considered negative.

Cosine similarity was computed for all intra-sequence and cross-sequence pairings. Nevertheless, only positive-positive and positive-negative pairs were simultaneously employed for model optimization according to the normalized temperature-scaled cross-entropy loss function Chen et al. (2020):

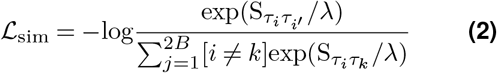

where S_*ik*_ denotes the cosine similarity between two normalized metric embeddings according to Equation 1, [*i* ≠ *k*] evaluates to 1 if *i* ≠ *k* and 0 otherwise, and *λ* is a scaling hyperparameter.

The loss minimizes the distance between embeddings of positive-positive pairs while maximizing the distances between embeddings of positive-negative pairs in the batch. Consequently, local representations *H*_*τ*_ of the underlying signal source are extracted while the loss is expected to suppresses other superimposed sources and noise.

### Probability map estimation

An independent MLP projection network *z*_*c*_ followed by sigmoid activation were employed to reduce the rank of the top-most embeddings transforming *H*_1_ into probability distribution *Ŷ* ∈ R^1^ of the P wave occurrence:

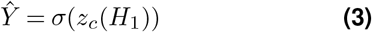

The function *z*_*c*_ was trained by employing same strategy as for the time-contrastive learning approach. Probability distribution *Ŷ* was by divided into a bag of temporal probabilities indexed by *τ* = 1, …, T. A ground truth label 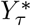 was assigned to each bag *Ŷ* _*τ*_ according to MIL paradigm denoted previously. To learn multiple instances, temporal pooling was applied to aggregate probabilities for each *Ŷ* _*τ*_. Here we continue with the maximum pooling (Boyd and Vandenberghe, 2004), but other choices (Zhou and Zhang, 2002; Ilse et al., 2018) are generally possible:

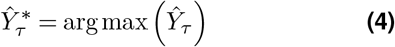

where 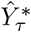 denotes the probability of the bag *τ* containing an instance of the P wave. The probability distribution was optimized using the weighted cross-entropy loss:

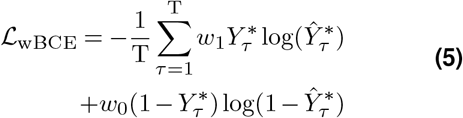

where *w*_1_, *w*_0_ balances the contribution of positive and negative class to the overall loss based on the inverse of its frequency of occurrence.

### Training setup

The AdamW algorithm (Kingma and Ba, 2015) with de-coupled L_2_ regularization (Loshchilov and Hutter, 2019) and default *β*_1_, *β*_2_, as recommended by Kingma and Ba (2015), was employed for the optimization. Convolutional and normalization layers were initialized using the Kaiming (He et al., 2015) and constant initialization, respectively. The initial learning rate *μ*_0_ = 0.001, batch size of 64, and temperature scaling factor *λ* = 0.15 were determined through the grid search. Learning rate was scheduled using warm-up phase for the first 20 epochs, followed by a reduce-on-plateau strategy with decaying factor of 0.1. The code was implemented in Python 3.8.1 (Van Rossum and Drake Jr, 1995) using the PyTorch 1.12.1 (Paszke et al., 2019). The model training was performed on the NVIDIA RTX 3060 12 G graphics card.

## Results and discussion

### Evaluation

To assess the performance of the proposed method (S-TCRL) and baseline models, we employed several metrics. The Dice score (DS_MIL_) was computed to evaluate the overlap between the model’s predicted probability of P-wave presence 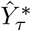 and the corresponding ground truth labels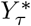. A threshold of 0.5 was applied to the P-wave probability scores *Ŷ* to convert them into binary classifications for recall and precision calculations of predicted fiducial points: P wave onset (P_on_), offset (P_off_), and peak (P_peak_). Due to limitations in the ground truth data regarding the exact location of each fiducial point, a tolerance of 60 ms was used when estimating both recall and precision.

### Experiments

The proposed S-TCRL method was compared to established baselines for weakly supervised classification using two different MIL-based deep learning approaches, described previously in Hejc et al. (2023). Both approaches utilized the same ground truth labels 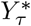. In the first approach (L-MIL), training occurred only through the final classification layer. In the second approach (FPN-MIL), each decoding layer of the FPN was trained independently. L-MIL essentially corresponds to fine-tuning the S-TCRL probability estimator introduced in Section. Penalization of each MIL learner was implemented using Equation 5.

Table 1 summarizes the performance of S-TCRL compared to L-MIL and FPN-MIL baselines on the entire validation dataset (DS_V_) and the sinus rhythm subgroup (DS_V_:SR). S-TCRL outperformed both baselines in P-wave detection, achieving a DS_MIL_ of 81.9% compared to 81.1% and 80.1% for L-MIL and FPN-MIL, respectively.

**Table 1.**
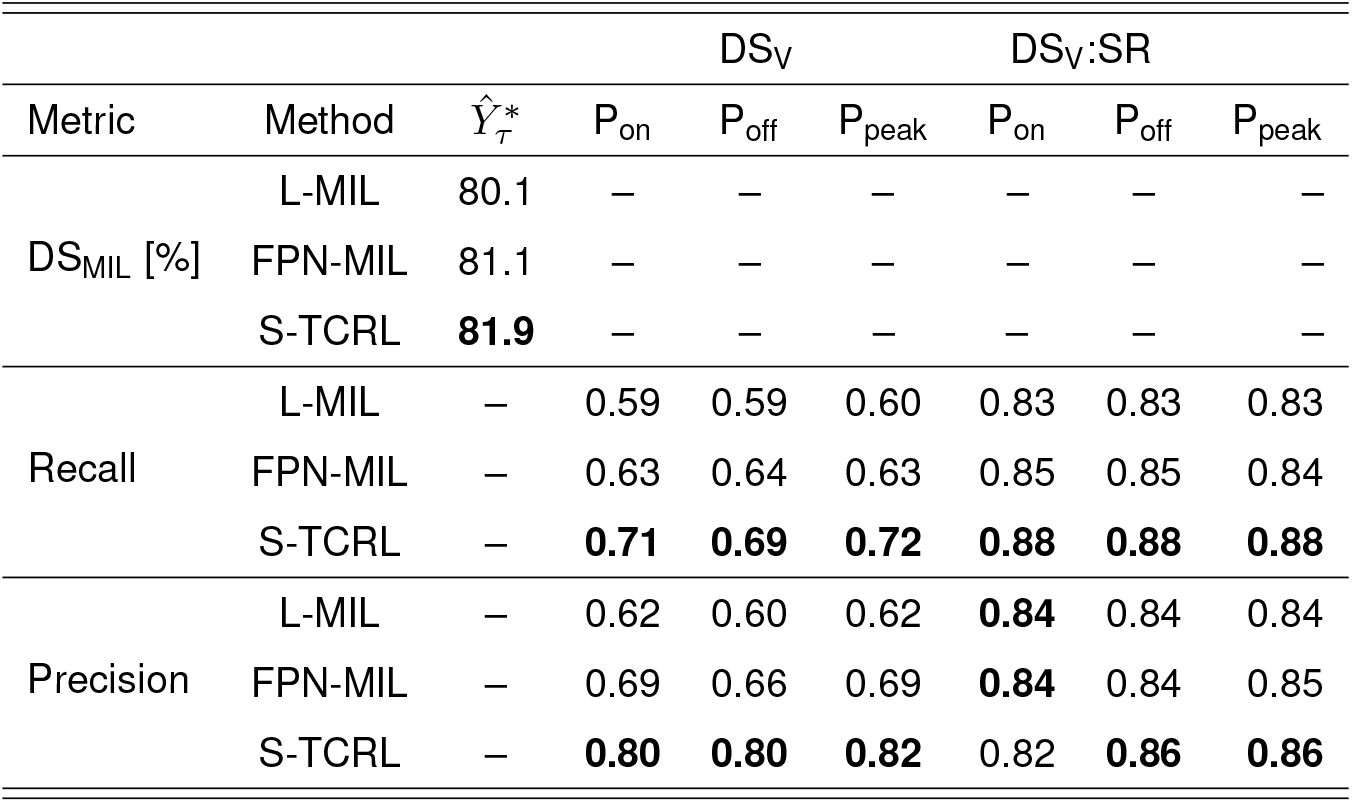
The performance of the S-TCRL, L-MIL, and FPN-MIL models on the validation set DS_V_ and sinus rhythm subset DS_V_:SR.

The proposed S-TCRL method demonstrates significant improvement over both baseline approaches. Compared to L-MIL, S-TCRL achieves a substantial improvement in P_peak_ detection, with recall increasing by 0.12 and precision by 0.20. Similarly, compared to FPN-MIL, S-TCRL shows improvement in recall (0.09) and precision (0.13). These results strongly suggest that contrastive learning plays a crucial role in enhancing the quality of P wave feature extraction.

The performance of all three methods in detecting key points of P waves within the DS_V_ subgroup exhibited consistency, with recall and precision exceeding 0.80 for all key points. This subgroup encompasses both well-defined P waves and less severe cardiac rhythm disorders, such as premature atrial contractions, ventricular extrasystoles, sinus tachycardia, or first-degree AV node blocks. These findings align with observations from Saclova et al. (2022), who reported significant variations in P wave detection performance across different databases. Their study found that recall and precision ranged from 78.1-93.1% and 72.0-88.6%, depending on the specific pathologies included in the dataset.

### Probability maps

Figure 3 illustrates P wave probability maps extracted by three methods: S-TCRL, L-MIL, and FPN-MIL. These maps indicate the likelihood of a P wave being present at each point in the ECG signal.

**Figure 3.**
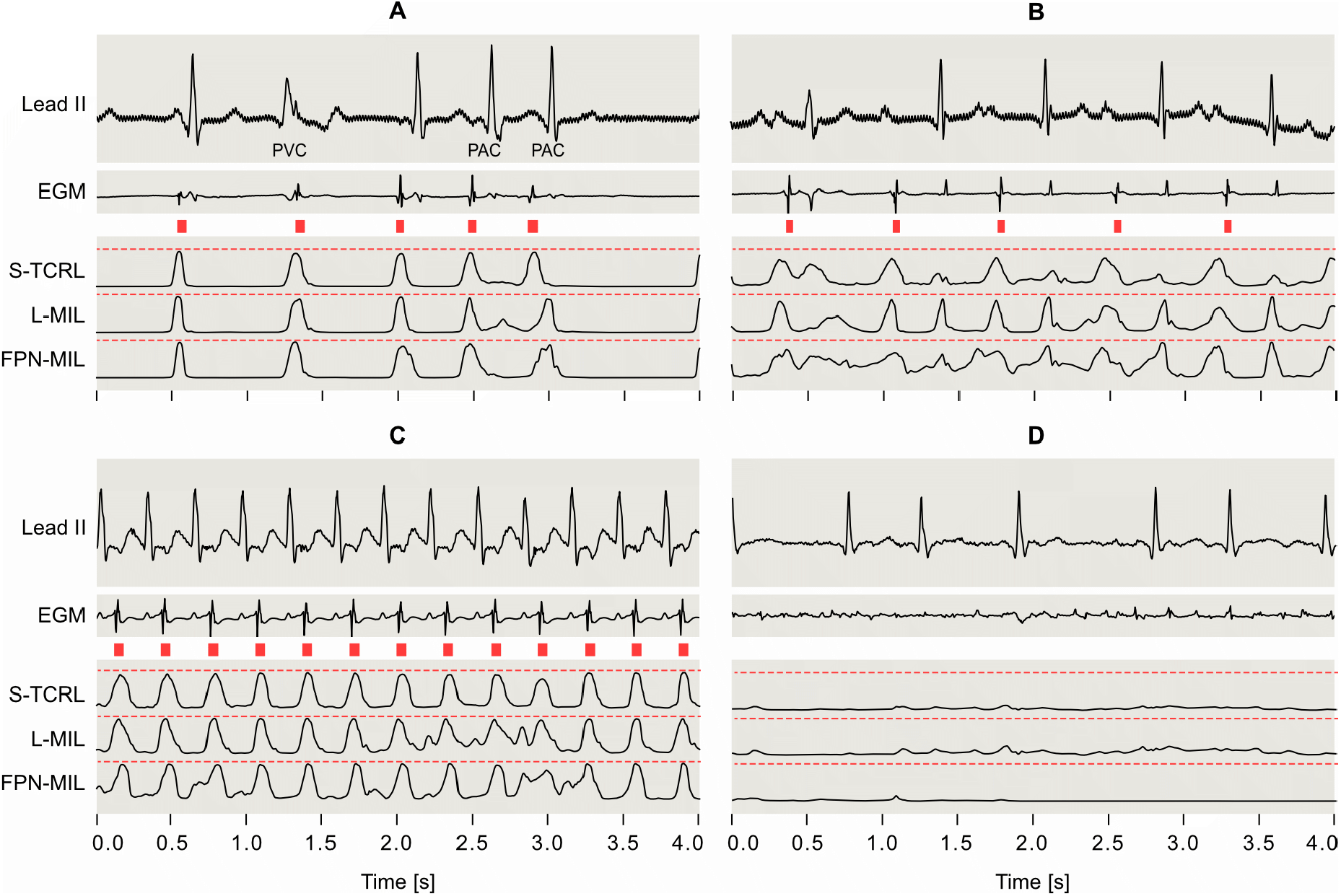
Representative ECG signals with P wave probability maps extracted by S-TCRL, L-MIL, and FPN-MIL methods for recordings containing: (A) multiple premature atrial (PAC) and ventricular (PVC) contractions, (B) sinus rhythm with atrioventricular block, (C) atrioventricular nodal re-entry tachycardia, and (D) atrial fibrillation. Top panels show the Lead II of the original ECG signals. Red rectangles in the middle denote surrogate labels for P waves. EGM refers to the intracardiac electrogram used to derive the surrogate labels.

Panels 3-A and 3-B demonstrate the models’ ability to extract P waves in irregular rhythms caused by premature beats (PACs) and atrioventricular block, where P waves were obscured by ventricular activity.

Panel 3-C showcases extraction in a signal with narrow QRS supraventricular tachycardia, where retrogradely activated P waves are embedded within the QRS complex and T waves, often challenging for automated segmentation methods but sometimes identifiable by experienced cardiologists.

Panel 3-D depicts model behavior on atrial fibrillation (AF) signals. The characteristic highly irregular activity of AF is evident, especially in the intracardiac EGM. The corresponding feature maps exhibit minimal fluctuations, reflecting the models’ ability to correctly identify the absence of regular P waves despite atrial activity potentially resembling P waves. AF classification is a widely studied topic due to associated complication risks. While conventional deep learning algorithms for AF classification struggle with interpreting extracted features, the proposed method offers clear and clinically interpretable parameters.

### Limitations

P wave segmentation in the presence of arrhythmias is a non-trivial task, as demonstrated by our results. Conventional weakly-supervised learning employing the MIL paradigm may not be sufficient for the effective extraction of microfeatures representing superimposed waves. Visual analysis showed that the models in some cases tended to approximate the temporal co-occurrence of the P-QRS-T waves, resulting in false positive detections. This is likely due to several factors: a) the complexity of the optimization goal and insufficient training data for the model to learn uncorrelated representations of superimposed wave; b) suboptimal settings for temporal hyperparameters based on prior knowledge about expected wave durations.

Novel regularization techniques might be necessary to address potential overfitting to the P-QRS-T temporal distribution, especially in deep learning architectures. A comparative study by Saclova et al. (2022); Maršánová et al. (2019) demonstrated that machine learning algorithms often overestimate performance on readily identifiable P waves. Deep learning models are likely susceptible to a similar effect. Therefore, ECG segmentation algorithms should be rigorously evaluated using diverse cardiac rhythm datasets.

Intracardiac EGMs have been used in only one prior study for P wave detection by Lenis et al. (2016). They employed a multi-electrode basket catheter to define reference positions for atrial activation. Basket catheters capture a broader spatiotemporal voltage distribution compared to the coronary sinus recordings used here, potentially providing a less incomplete positional reference. However, basket catheters are not routinely used in EP studies, hindering their applicability for constructing large training datasets.

## Conclusion

The proposed S-TCRL method, leveraging surrogate labels and temporal contrastive learning, demonstrates promising potential for capturing the temporal dynamics of superimposed ECG waves, even with limited labeled data. Surrogate labels enable the extraction of more context-rich intrinsic representations compared to existing ECG processing methods, even in challenging arrhythmias like atrial fibrillation and supraventricular tachycardia. This makes S-TCRL a valuable tool for automated ECG interpretation and diagnosis.

Furthermore, S-TCRL stands out as the first work to successfully embed representations of hidden P waves into a deep learning model. Further research is warranted to validate S-TCRL in larger clinical studies and explore its potential integration into clinical decision-making support systems.

## Bibliography

Akhbari, M., Shamsollahi, M. B., Jutten, C., Armoundas, A. A., and Sayadi, O. (2016). ECG denoising and fiducial point extraction using an extended kalman filtering framework with linear and nonlinear phase observations. Physiological Measurement, 37(2):203–226. doi: 10.1088/0967-3334/37/2/203.

Bachman, P., Hjelm, R. D., and Buchwalter, W. Learning representations by maximizing mutual information across views. Curran Associates Inc., Red Hook, NY, USA, (2019).

Banville, H., Chehab, O., Hyvärinen, A., Engemann, D.-A., and Gramfort, A. (2021). Uncovering the structure of clinical eeg signals with self-supervised learning. Journal of Neural Engineering, 18(4):046020. doi: 10.1088/1741-2552/abca18.

Boyd, S. and Vandenberghe, L. Convex Optimization. Cambridge University Press, (2004). doi: 10.1017/cbo9780511804441. URL 10.1017/cbo9780511804441.

Brady, W. J., Mattu, A., Tabas, J., and Ferguson, J. D. (2017). The differential diagnosis of wide QRS complex tachycardia. The American Journal of Emergency Medicine, 35(10): 1525–1529. doi: 10.1016/j.ajem.2017.07.056.

Brugada, P., Brugada, J., Mont, L., Smeets, J., and Andries, E. W. (1991). A new approach to the differential diagnosis of a regular tachycardia with a wide QRS complex. Circulation, 83(5):1649–1659. doi: 10.1161/01.cir.83.5.1649.

Chen, L. Y., Chung, M. K., Allen, L. A., Ezekowitz, M., Furie, K. L., McCabe, P., Noseworthy, P. A., Perez, M. V., and Turakhia, M. P. (2018). Atrial fibrillation burden: Moving beyond atrial fibrillation as a binary entity: A scientific statement from the american heart association. Circulation, 137(20). doi: 10.1161/cir.0000000000000568.

Chen, T., Kornblith, S., Norouzi, M., and Hinton, G. E. (2020). A simple framework for contrastive learning of visual representations. CoRR, abs/2002.05709.

Cheng, J. Y., Goh, H., Dogrusoz, K., Tuzel, O., and Azemi, E. Subject-aware contrastive learning for biosignals, (2020).

Costandy, R. N., Gasser, S. M., El-Mahallawy, M. S., Fakhr, M. W., and Marzouk, S. Y. (2020). P-wave detection using a fully convolutional neural network in electrocardiogram images. Applied Sciences, 10(3):976. doi: 10.3390/app10030976.

Dave, I., Gupta, R., Rizve, M. N., and Shah, M. (2022). TCLR: Temporal contrastive learning for video representation. Computer Vision and Image Understanding, 219:103406. doi: 10.1016/j.cviu.2022.103406.

Di Marco, L. Y. and Chiari, L. (2011). A wavelet-based ECG delineation algorithm for 32-bit integer online processing. Biomed. Eng. Online, 10(1):23.

Duraj, K., Piaseczna, N., Kostka, P., and Tkacz, E. (2022). Semantic segmentation of 12-lead ECG using 1d residual u-net with squeeze-excitation blocks. Applied Sciences, 12 (7):3332. doi: 10.3390/app12073332.

Gonthier, N., Ladjal, S., and Gousseau, Y. (2022). Multiple instance learning on deep features for weakly supervised object detection with extreme domain shifts. Computer Vision and Image Understanding, 214:103299. doi: 10.1016/j.cviu.2021.103299.

Gutmann, M. and Hyvärinen, A. Noise-contrastive estimation: Anew estimation principle for unnormalized statistical models. In Teh, Y. W.and Titterington, M., editors, Proceedings of the Thirteenth International Conference on Artificial Intelligence and Statistics, volume 9 of Proceedings of Machine Learning Research, pages 297–304, Chia Laguna Resort, Sardinia, Italy, (2010). PMLR. URL https://proceedings.mlr.press/v9/gutmann10a.html.

Hadsell, R., Chopra, S., and LeCun, Y. Dimensionality reduction by learning an invariant mapping. In 2006 IEEE Computer Society Conference on Computer Vision and Pattern Recognition (CVPR’06), volume 2, pages 1735–1742, (2006). doi: 10.1109/CVPR.2006.100.

He, K., Zhang, X., Ren, S., and Sun, J. Delving Deep into Rectifiers: Surpassing Human-Level Performance on ImageNet Classification. In 2015 IEEE International Conference on Computer Vision (ICCV), pages 1026–1034, (2015). doi: 10.1109/ICCV.2015.123.

Hejc, J., Redina, R., Pospisil, D., Rakova, I., Kolarova, J., and Starek, Z. Weakly supervised p wave segmentation in pathological electrocardiogram signals using deep multiple| 8 instance learning. In 2023 Computing in Cardiology (CinC), volume 50, pages 1–4, (2023). doi: 10.22489/CinC.2023.321.

Hejc, J., Redina, R., Kolarova, J., and Starek, Z. (2024). Multi-channel delineation of intracardiac electrograms for arrhythmia substrate analysis using implicitly regularized convolutional neural network with wide receptive field. Biomedical Signal Processing and Control, 94:106274. doi: 10.1016/j.bspc.2024.106274.

Hénaff, O. J., Srinivas, A., De Fauw, J., Razavi, A., Doersch, C., Eslami, S. M. A., and Van Den Oord, A. Data-efficient image recognition with contrastive predictive coding. In Proceedings of the 37th International Conference on Machine Learning, ICML’20. JMLR.org, (2020).

Hernández, A. I., Carrault, G., and Mora, F. (2000). Improvement of a p-wave detector by a bivariate classification stage. Transactions of the Institute of Measurement and Control, 22(3):231–242. doi: 10.1177/014233120002200303.

Hesar, H. D. and Mohebbi, M. (2019). A multi rate marginalized particle extended kalman filter for p and t wave segmentation in ecg signals. IEEE Journal of Biomedical and Health Informatics, 23(1):112–122. doi: 10.1109/JBHI.2018.2794362.

Hossain, M. B., Bashar, S. K., Walkey, A. J., McManus, D. D., and Chon, K. H. (2019). An accurate qrs complex and p wave detection in ecg signals using complete ensemble empirical mode decomposition with adaptive noise approach. IEEE Access, 7:128869–128880. doi: 10.1109/ACCESS.2019.2939943.

Hyvarinen, A. and Morioka, H. Unsupervised feature extraction by time-contrastive learning and nonlinear ica. In Lee, D., Sugiyama, M., Luxburg, U., Guyon, I., and Garnett, R., editors, Advances in Neural Information Processing Systems, volume 29. Curran Associates, Inc., (2016). URL https://proceedings.neurips.cc/paper_files/paper/2016/file/d305281faf947ca7acade9ad5c8c818c-Paper.pdf.

Ilse, M., Tomczak, J., and Welling, M. Attention-based deep multiple instance learning. In International conference on machine learning, pages 2127–2136. PMLR, (2018).

Issa, Z. F., Miller, J. M., and Zipes, D. P. Electrophysiological Mechanisms of Cardiac Arrhythmias, page 51–80. Elsevier, (2019). doi: 10.1016/b978-0-323-52356-1.00003-7. URL 10.1016/b978-0-323-52356-1.00003-7.

Katritsis, D. G., Boriani, G., Cosio, F. G., Hindricks, G., Jaïs, P., Josephson, M. E., Keegan, R., Kim, Y.-H., Knight, B. P., Kuck, K.-H., Lane, D. A., Lip, G. Y. H., Malmborg, H., Oral, H., Pappone, C., Themistoclakis, S., Wood, K. A., Blomström-Lundqvist, C., Gorenek, B., Dagres, N., Dan, G.-A., Vos, M. A., Kudaiberdieva, G., Crijns, H., Roberts-Thomson, K., Lin, Y.-J., Vanegas, D., Caorsi, W. R., Cronin, E., and Rickard, J. (2016). European heart rhythm association (ehra) consensus document on the management of supraventricular arrhythmias, endorsed by heart rhythm society (hrs), asia-pacific heart rhythm society (aphrs), and sociedad latinoamericana de estimulación cardiaca y electrofisiologia (solaece). EP Europace, 19(3):465–511. doi: 10.1093/europace/euw301.

Kimura-Medorima, S. T., Lino, A. P. B. L., Almeida, M. P. C., Figueiredo, M. J. O., da Mota Silveira-Filho, L., de Oliveira, P. P. M., Coelho, O. R., Souza, J. R. M., Nadruz, W., Petrucci, O., and Sposito, A. C. (2018). P-wave duration is a predictor for long-term mortality in post-CABG patients. PLOS ONE, 13(7):e0199718. doi: 10.1371/journal.pone.0199718.

Kingma, D. P. and Ba, J. Adam: A method for stochastic optimization. In Bengio, Y.and LeCun, Y., editors, 3rd International Conference on Learning Representations, ICLR 2015, San Diego, CA, USA, May 7-9, 2015, Conference Track Proceedings, (2015). URL http://arxiv.org/abs/1412.6980.

Kiyasseh, D., Zhu, T., and Clifton, D. A. Clocs: Contrastive learning of cardiac signals across space, time, and patients. In Meila, M.and Zhang, T., editors, Proceedings of the 38th International Conference on Machine Learning, volume 139 of Proceedings of Machine Learning Research, pages 5606–5615. PMLR, (2021). URL https://proceedings.mlr.press/v139/kiyasseh21a.html.

Lenis, G., Pilia, N., Oesterlein, T., Luik, A., Schmitt, C., and Dössel, O. (2016). P wave detection and delineation in the ECG based on the phase free stationary wavelet transform and using intracardiac atrial electrograms as reference. Biomedical Engineering / Biomedizinische Technik, 61(1):37–56. doi: 10.1515/bmt-2014-0161.

Li, C., Zheng, C., and Tai, C. (1995). Detection of ECG characteristic points using wavelet transforms. IEEE Transactions on Biomedical Engineering, 42(1):21–28. doi: 10.1109/10.362922.

Lin, T.-Y., Dollar, P., Girshick, R., He, K., Hariharan, B., and Belongie, S. Feature pyramid networks for object detection. In Proceedings of the IEEE Conference on Computer Vision and Pattern Recognition (CVPR), (2017).

Liu, Z., Alavi, A., Li, M., and Zhang, X. (2023). Self-supervised contrastive learning for medical time series: A systematic review. Sensors, 23(9):4221. doi: 10.3390/s23094221.

Loshchilov, I. and Hutter, F. Decoupled weight decay regularization. In International Conference on Learning Representations, (2019). URL https://openreview.net/forum?id=Bkg6RiCqY7.

Malali, A., Hiriyannaiah, S., G.M., S., K.G., S., and N.T., S. (2020). Supervised ECG wave segmentation using convolutional LSTM. ICT Express, 6(3):166–169. doi: 10.1016/j.icte.2020.04.004.

Marenco, J. P., Nakagawa, H., Yang, S., MacAdam, D., Xu, L., He, D. S., Link, M. S., Homoud, M. K., III, N. M. E., and Wang, P. J. (2003). Testing of a new t-wave subtraction algorithm as an aid to localizing ectopic atrial beats. Annals of Noninvasive Electrocardiology, 8(1): 55–59. doi: 10.1046/j.1542-474x.2003.08109.x.

Maršánová, L., Němcová, A., Smíšek, R., Vítek, M., and Smital, L. (2019). Advanced p wave detection in ecg signals during pathology: Evaluation in different arrhythmia contexts. Scientific Reports, 9(1). doi: 10.1038/s41598-019-55323-3.

Martínez, A., Alcaraz, R., and Rieta, J. J. (2010). Application of the phasor transform for automatic delineation of single-lead ECG fiducial points. Physiological Measurement, 31 (11):1467–1485. doi: 10.1088/0967-3334/31/11/005.

Martinez, J., Almeida, R., Olmos, S., Rocha, A., and Laguna, P. (2004). A wavelet-based ecg delineator: evaluation on standard databases. IEEE Transactions on Biomedical Engineering, 51(4):570–581. doi: 10.1109/TBME.2003.821031.

McSharry, P., Clifford, G., Tarassenko, L., and Smith, L. (2003). A dynamical model for generating synthetic electrocardiogram signals. IEEE Transactions on Biomedical Engineering, 50(3):289–294. doi: 10.1109/TBME.2003.808805.

Mehari, T. and Strodthoff, N. (2022). Self-supervised representation learning from 12-lead ecg data. Computers in Biology and Medicine, 141:105114. doi: 10.1016/j.compbiomed.2021.105114.

Panigrahy, D. and Sahu, P. K. (2018). P and T wave detection and delineation of ECG signal using differential evolution (DE) optimization strategy. Australas. Phys. Eng. Sci. Med., 41(1):225–241.

Paszke, A., Gross, S., Massa, F., Lerer, A., Bradbury, J., Chanan, G., Killeen, T., Lin, Z., Gimelshein, N., Antiga, L., Desmaison, A., Kopf, A., Yang, E., DeVito, Z., Raison, M., Tejani, A., Chilamkurthy, S., Steiner, B., Fang, L., Bai, J., and Chintala, S. Pytorch: An imperative style, high-performance deep learning library. In Advances in Neural Information Processing Systems 32, pages 8024–8035. Curran Associates, Inc., (2019). URL http://papers.neurips.cc/paper/9015-pytorch-an-imperative-style-high-performance-deep-learning-library.pdf.

Peimankar, A. and Puthusserypady, S. An ensemble of deep recurrent neural networks for pwave detection in electrocardiogram. In ICASSP 2019 – 2019 IEEE International Conference on Acoustics, Speech and Signal Processing (ICASSP), pages 1284–1288, (2019). doi: 10.1109/ICASSP.2019.8682307.

Portet, F. (2008). P wave detector with PP rhythm tracking: evaluation in different arrhythmia contexts. Physiol. Meas., 29(1):141–155.

Qi, J., Shi, P., Hu, L., Zhang, T., and Xie, S. (2019). ECG characteristic wave detection based on deep recursive long short-term memory. Journal of Medical Imaging and Health Informatics, 9(9):1920–1924. doi: 10.1166/jmihi.2019.2815.

Rieta, J., Castells, F., Sanchez, C., Zarzoso, V., and Millet, J. (2004). Atrial activity extraction for atrial fibrillation analysis using blind source separation. IEEE Transactions on Biomedical Engineering, 51(7):1176–1186. doi: 10.1109/tbme.2004.827272.

Roberts-Thomson, K. C., Kistler, P. M., and Kalman, J. M. (2006). Focal atrial tachycardia i: Clinical features, diagnosis, mechanisms, and anatomic location. Pacing and Clinical Electrophysiology, 29(6):643–652. doi: 10.1111/j.1540-8159.2006.00413.x.

Saclova, L., Nemcova, A., Smisek, R., Smital, L., Vitek, M., and Ronzhina, M. (2022). Reliable P wave detection in pathological ECG signals. Sci. Rep., 12(1):6589.

Sahambi, J., Tandon, S., and Bhatt, R. (1997). Using wavelet transforms for ECG characterization. an on-line digital signal processing system. IEEE Engineering in Medicine and Biology Magazine, 16(1):77–83. doi: 10.1109/51.566158.

Sarkar, P. and Etemad, A. (2022). Self-supervised ecg representation learning for emotion recognition. IEEE Transactions on Affective Computing, 13(3):1541–1554. doi: 10.1109/TAFFC.2020.3014842.

Schroff, F., Kalenichenko, D., and Philbin, J. Facenet: A unified embedding for face recognition and clustering. In 2015 IEEE Conference on Computer Vision and Pattern Recognition (CVPR), pages 815–823, (2015). doi: 10.1109/CVPR.2015.7298682.

Sermanet, P., Lynch, C., Hsu, J., and Levine, S. Time-contrastive networks: Self-supervised learning from multi-view observation. In 2017 IEEE Conference on Computer Vision and Pattern Recognition Workshops (CVPRW), pages 486–487, (2017). doi: 10.1109/CVPRW.2017.69.

Sippensgroe-newegen, A., Mlynash, M. D., Roithinger, F. X., Goseki, Y., and Lesh, M. D. (2001). Electrocardiographic analysis of ectopic atrial activity obscured by ventricular repolarization: P wave isolation using an automatic 62-lead QRST subtraction algorithm. Journal of Cardiovascular Electrophysiology, 12(7):780–790. doi: 10.1046/j.1540-8167.2001.00780.x.

Spicher, N. and Kukuk, M. (2020). Delineation of electrocardiograms using multiscale parameter estimation. IEEE Journal of Biomedical and Health Informatics, 24(8):2216–2229. doi: 10.1109/JBHI.2019.2963786.

Stafford, P. J., Turner, I., and Vincent, R. (1991). Quantitative analysis of signal-averaged p waves in idiopathic paroxysmal atrial fibrillation. The American Journal of Cardiology, 68 (8):751–755. doi: 10.1016/0002-9149(91)90648-5.

Stridh, M. and Sommo, L. (2001). Spatiotemporal QRST cancellation techniques for analysis of atrial fibrillation. IEEE Transactions on Biomedical Engineering, 48(1):105–111. doi: 10.1109/10.900266.

Tereshchenko, L. G., Waks, J. W., Kabir, M., Ghafoori, E., Shvilkin, A., and Josephson, M. E. (2015). Analysis of speed, curvature, planarity and frequency characteristics of heart vector movement to evaluate the electrophysiological substrate associated with ventricular tachycardia. Computers in Biology and Medicine, 65:150–160. doi: 10.1016/j.compbiomed.2015.03.001.

van den Oord, A., Li, Y., and Vinyals, O. Representation learning with contrastive predictive coding, (2019).

Van Rossum, G. and Drake Jr, F. L. Python reference manual. Centrum voor Wiskunde en Informatica Amsterdam, (1995).

Vaya, C., Rieta, J., Sanchez, C., and Moratal, D. (2007). Convolutive blind source separation algorithms applied to the electrocardiogram of atrial fibrillation: Study of performance. IEEE Transactions on Biomedical Engineering, 54(8):1530–1533. doi: 10.1109/tbme.2006.889778.

Vicar, T., Hejc, J., Novotna, P., Ronzhina, M., and Janousek, O. Ecg abnormalities recognition using convolutional network with global skip connections and custom loss function. In 2020 Computing in Cardiology, pages 1–4, (2020). doi: 10.22489/CinC.2020.189.

Vicar, T., Novotna, P., Hejc, J., Janousek, O., and Ronzhina, M. Cardiac abnormalities recognition in ecg using a convolutional network with attention and input with an adaptable number of leads. In 2021 Computing in Cardiology (CinC), volume 48, pages 1–4, (2021). doi: 10.23919/CinC53138.2021.9662806.

Yuan, Y., Xun, G., Suo, Q., Jia, K., and Zhang, A. (2019). Wave2vec: Deep representation learning for clinical temporal data. Neurocomputing, 324:31–42. doi: 10.1016/j.neucom.2018.03.074. Deep Learning for Biological/Clinical Data.

Zhou, Z.-H. and Zhang, M.-L. Neural networks for multi-instance learning. In Proceedings of the International Conference on Intelligent Information Technology, Beijing, China, pages 455–459, (2002).

